# Efficient Optimization of Genotype Pairs for Intercropping using Genomic Prediction and Bayesian Optimization

**DOI:** 10.64898/2026.05.15.725387

**Authors:** Sei Kinoshita, Hiroyoshi Iwata

## Abstract

Intercropping is a promising strategy to improve productivity and sustainability in agricultural systems, but designing effective genotype combinations remains a major challenge owing to the rapid increase in possible pairings as the number of candidate genotypes increases. This creates a practical bottleneck because field evaluation of all combinations is infeasible under realistic resource constraints. Here, we propose a framework that integrates genomic prediction and Bayesian optimization to support efficient decision-making for intercropping system design. Using genome-wide marker data from sorghum and soybean, we simulated intercropping performance across 5,214 genotype pairs under certain genetic architectures, including variation in heritability, correlations between direct and indirect genetic effects, and the contribution of pair-specific interactions. Genomic prediction models incorporating direct and indirect genetic effects substantially improved prediction accuracy compared with models based on direct genetic effects alone, and inclusion of specific mixing ability further enhanced the performance under high-heritability conditions. When coupled with Bayesian optimization, the models rapidly identified superior genotype pairs, requiring fewer evaluation cycles than random or prediction-only search strategies. Acquisition functions that account for predicted uncertainty were most effective in complex scenarios involving interaction effects or negative correlations between direct and indirect effects. These results demonstrate that combining genomic prediction with Bayesian optimization can substantially reduce the experimental burden associated with intercropping design, while improving the efficiency of identifying high-performing genotype pairs. The proposed framework provides a practical approach for prioritizing candidate mixtures in breeding and field evaluation, and contributes to the development of data-driven strategies for sustainable agricultural systems.

**Highlights:**

- A data-driven framework was developed to optimize genotype pairs in intercropping.
- Modeling indirect effects improved prediction accuracy across genotype pairs.
- Pair-specific interactions enhanced prediction under high-heritability conditions.
- Bayesian optimization identified superior pairs under limited evaluation capacity.
- The framework reduces field-testing requirements for intercropping system design.

## 1. Introduction

Modern agriculture has achieved substantial gains in crop productivity through the development of high-yielding cultivars and the widespread use of synthetic fertilizers, pesticides, and mechanization. However, although these advances have greatly increased food production, they have also imposed severe environmental costs, including soil degradation, deterioration of water quality, and loss of biodiversity (Jing et al., 2025). As a result, there is growing interest in cropping systems that can maintain productivity while reducing external inputs and ecological impacts. In this context, intercropping, defined as the simultaneous cultivation of two or more crop species in the same field, has received renewed attention as an alternative to monoculture (Li et al., 2023; Toker et al., 2024). Intercropping is a traditional agricultural strategy that has been used for centuries in many regions of the world (Harwood, 2024). A growing body of evidence indicates that intercropping can enhance overall productivity and land-use efficiency, improve yield stability under variable environmental conditions, and contribute to pest and disease suppression through increased on-farm biodiversity and associated ecological interactions (Akchaya et al., 2025).

Despite these advantages, the practical design of intercropping systems remains challenging. Most contemporary cultivars have been developed for monoculture systems and their performance is not necessarily maintained under intercropping (Dubey et al., 2024). Therefore, identifying combinations of genotypes that interact favorably is essential to improve the productivity and stability of intercropping systems. While some researchers are focused on breeding cultivars specifically adapted to intercropping through crossing and selection (Moore et al., 2022; Abou Khater et al., 2024), the present study proposes a method to identify genotype pairs suited to intercropping from among existing cultivars and genetic resources prior to the initiation of such breeding programs.

Because many target traits in intercropping, including yield, are quantitative traits, the selection of suitable pairs of genotypes can be effectively approached within a quantitative genetic framework, particularly by modeling and evaluating the genetic potential of genotype pairs using genomic prediction based on genome-wide marker information (Annicchiarico et al., 2021; Bančič et al., 2021). In quantitative genetic models for intercropping, following the removal of the residual error from the phenotype, the genetic variation can be decomposed into 1) direct genetic effects (DGE), representing the effects of each crop genotype on its own performance, 2) indirect genetic effects (IGE), representing the effects of a genotype on its intercropped partner, and 3) specific mixing ability (SMA), representing genetic interactions that are expressed only in particular genotype pairs (Bijma, 2014; Forst et al., 2019; Bourke et al., 2021). DGE and IGE represent the average effects of a genotype across pairs and are also collectively termed the general mixing ability (GMA). Under this framework, genotype assessment should be conducted on a pairwise basis. However, as the number of candidate genotypes increases, the number of possible intercropping pairs grows explosively. For example, if 50 candidate genotypes are available for each crop, the total number of possible pairs reaches 2,500, a number impractical to evaluate in field trials. In practice, only a limited number of genotype pairs can be assessed in each season owing to time, labor, and cost constraints. Therefore, efficient methods are needed to identify suitable pairs from a vast number of candidates under such constraints.

Bayesian optimization is a machine-learning approach for efficiently finding the maximum or minimum of an objective function that is costly to evaluate in terms of experimental time or computational resources, while requiring only a limited number of evaluations. Bayesian optimization has the advantage of explicitly incorporating predictive uncertainty. By appropriately exploring unknown regions where the current model remains uncertain, rather than becoming trapped in local optima, it enables a more effective search for the global optimum (Siska et al., 2026). Recently, Bayesian optimization has been adopted in plant breeding, for example, to identify superior genotypes in a pre-breeding population by genomic selection and to optimize parameters in breeding schemes (Tanaka and Iwata, 2018; Diot and Iwata, 2023; Tu and Liao, 2026).

In the context of intercropping design, the objective function corresponds to the genotypic value of a genotype pair, as estimated from a genomic prediction model trained on phenotypic data obtained from field trials. Given the impracticality of evaluating all possible pairs directly in the field, this problem can be framed as the efficient identification of the maximum (i.e., an intercropping pair with the highest genotypic value) of an expensive-to-evaluate objective function under limited evaluation capacity (i.e., field trials). Thus, the search for superior intercropping pairs constitutes a decision-making problem that is particularly well suited to Bayesian optimization.

In this study, we propose a framework that integrates genomic prediction and Bayesian optimization to achieve the efficient design of intercropping systems. Using simulated phenotypic data representing diverse genetic architectures, we first evaluated the prediction accuracy of genomic prediction models that differ in their treatment of DGE, IGE, and SMA. We then coupled an appropriate genomic prediction model with Bayesian optimization and assessed its performance in identifying superior genotype pairs, in comparison with random search and prediction-based selection. Our objective was to demonstrate how data-driven methods can improve decision-making and reduce the experimental burden in the design of intercropping systems.

## 2. Materials and Methods

### 2.1. Genotype data

Marker genotype data were obtained from publicly available sorghum and soybean genetic resources. For sorghum, we selected 237 accessions from the 289-accession sorghum germplasm panel compiled by Takanashi et al. (2022). The original panel comprised accessions obtained from public bioresources, including the ICRISAT sorghum collection, NARO Genebank, and the USDA germplasm collection. For soybean, we used the whole-genome sequencing data for 198 accessions obtained by Kajiya-Kanegae et al. (2021). Of these 198 accessions, 192 were derived from Japanese and global soybean mini-core collections (Kaga et al., 2012). To reduce the computational cost of simulations, the number of sorghum and soybean accessions was decreased to one-third for each species. Specifically, pairwise Euclidean distances were calculated from the genotype matrix and representative accessions were selected using the *k*-medoids clustering algorithm. As a result, 79 sorghum accessions and 66 soybean accessions were chosen for subsequent analyses. The relative distribution of the selected and non-selected accessions in the full panel was visualized in a principal component analysis (PCA) ordination based on the genomic relationship matrix (Figure S1).

In the genomic data for the original panels (237 sorghum and 198 soybean accessions), 3,393,214 and 4,776,813 single-nucleotide polymorphisms (SNPs), respectively, were identified. After filtering to retain only markers with a minor allele frequency (MAF) > 0.1, 1,891,947 SNPs remained for sorghum and 2,606,111 SNPs remained for soybean. To further reduce the computational burden, only markers with linkage disequilibrium (LD) < 0.6 were retained, resulting in 91,692 SNPs for sorghum and 61,426 SNPs for soybean. In addition, for the 79 sorghum accessions and 66 soybean accessions selected by *k*-medoids clustering, the SNPs were further filtered to retain markers with MAF > 0.1. The final sets of 87,535 and 57,717 SNPs for sorghum and soybean, respectively, were used for subsequent pseudo-phenotype generation.

### 2.2. Construction of the linkage map

Map distances were estimated from the physical positions of the SNP markers by converting the physical distance to genetic distance using species-specific average genome-wide recombination rates. For sorghum, a reported total genetic map length of 1603.5 cM and a genome size of 730 Mb (Mace et al., 2009; Paterson et al., 2009) were used to calculate the conversion rate (1603.5 cM/730 Mb = 2.2 cM/Mb), which was then applied to estimate the map distances. For soybean, an average genome-wide recombination rate of 2.5 cM/Mb was used based on previous studies (Lee et al., 2020).

### 2.3. Generation of pseudo-phenotype data

In intercropping systems, the phenotypic value can be decomposed into general mixing ability (GMA), specific mixing ability (SMA), and a residual term that includes environmental factors. GMA represents the average performance of a given genotype in intercropping and can be further divided into direct genetic effects (DGE) and indirect genetic effects (IGE). DGE refers to the influence of the focal genotype on its own phenotypic value, whereas IGE describes the impact of the neighboring genotype on the phenotypic value of the focal genotype. GMA is determined for each genotype and is heritable, whereas SMA denotes non-heritable interactions that arise only in specific genotype pairs (Forst et al., 2019; Haug et al., 2021).

In line with these concepts, pseudo-phenotypes were generated for each species for all 5,214 possible genotype pairs between the 79 sorghum accessions and the 66 soybean accessions. For notational convenience in the equations, sorghum and soybean are hereafter denoted as crop A and crop B, respectively. For the genotype pair consisting of crop A genotype *i* and crop B genotype *j*, the phenotypes were generated as

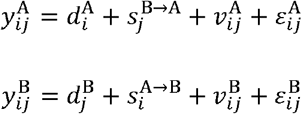

where 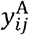 and 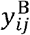 denote the phenotypic values of crop A and crop B, respectively, in genotype pair (*i,j*). The terms 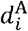 and 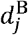 represent the DGE of genotypes *i* and *j* on their own phenotypes, respectively; 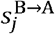 and 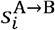 represent the IGE of the neighboring genotype on the focal genotype; and 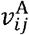 and 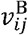 represent the SMA, which are pair-specific interaction effects that depend on the particular genotype pair. The residual terms are denoted by 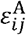 and 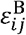. The equations for pseudo-phenotype generation can be expressed in matrix notation as follows:

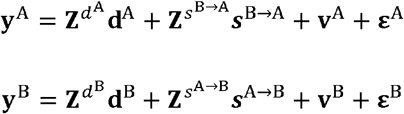

Where **y**^A^,**y**^B^,**y**^A^,**y**^B^, **ε**^A^ and **ε**^B^ are 5,214×1 vectors; **d**^A^ and ***s***^*A*→*B*^ are 79 ×1 × **d**^B^ and ***s***^*B*→*A*^are 66 × 1 vectors; and 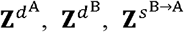, and 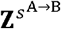 are design matrices.

Given that SMA is not heritable, heritability can be defined for crop A and B as follows:

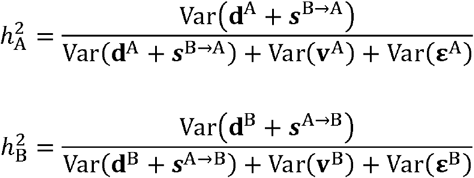

Because only additive genetic effects are considered for DGE and IGE, the heritability discussed here is also based solely on additive genetic variance.

To account for a range of phenotypic patterns observed in intercropping, we established a total of 18 pseudo-phenotype generation scenarios by considering the following three factors: heritability 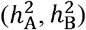, the genetic correlation between DGE and IGE, and the variance ratio of SMA to GMA. The heritability was set to 0.3 or 0.7, the genetic correlation between DGE and IGE was set to −0.7, −0.1, or 0.7, and the variance ratio of SMA to GMA was set to 0, 0.2, or 0.4. The rationale for introducing a genetic correlation between DGE and IGE is that, in intercropping systems, these effects are often reported to be correlated (Bourke et al., 2021). A negative correlation is typically interpreted as evidence of competition among neighboring genotypes, whereas a positive correlation is considered to reflect facilitation (Firmat and Litrico, 2022). In addition, we evaluated the scenario with a weak genetic correlation (−0.1). Furthermore, based on previous studies, SMA is often absent or accounts for a smaller proportion of the variance compared with GMA (Haug et al., 2023; Hohmann et al., 2026). Therefore, the proportion of variance attributed to SMA was set to be smaller than that of GMA.

Pseudo-phenotypes were generated using the following procedure.

1. The number of quantitative trait nucleotides (QTNs) was first determined. The positions and effects of the QTNs were then generated according to the genetic correlation between DGE and IGE. Specifically, 3,000 QTNs were assigned for sorghum (300 QTNs per chromosome) and 2,000 QTNs for soybean (100 QTNs per chromosome). A relatively large number of QTNs was used to approximate polygenic architectures. Following Neyhart et al. (2019), the positions and effects of the QTNs were generated such that the genetic correlation between DGE and IGE matched that specified for each scenario. Genetic correlations can arise through three mechanisms: pleiotropy, in which the QTNs for DGE and IGE are located at the same locus; tight linkage, in which they are located at distinct but closely linked loci; and loose linkage, in which they are located at more distant loci. In this study, genetic correlations were generated under a tight-linkage scenario. First, the positions of QTNs for DGE were sampled. For each QTN for DGE, a QTN for IGE was then randomly sampled from markers located within 5 cM of the corresponding QTN for DGE. If no marker was present within 5 cM, the nearest marker was selected when the QTN was at a chromosome end. When the QTN was not at the end of a chromosome, the marker with the higher LD was chosen from among the two nearest flanking markers.
2. DGE and IGE were then generated by combining the effects obtained in step 1 with the genotypic score matrix, and were subsequently scaled so that their variance components were equal (i.e., 1:1) to simplify the interpretation of their relative contributions.
3. SMA was sampled from a multivariate normal distribution with a variance–covariance structure given by the Kronecker product of **G**_A_ and **G**_B_ to model pairwise genetic interactions between genotypes of the two species. Here, **G**_A_ and **G**_B_ denote the additive genomic relationship matrix of sorghum and soybean, respectively.
4. SMA was then scaled according to the following equations so that the proportion of SMA variance relative to GMA corresponded to that specified for each scenario. Here, *γ*_A_ and *γ*_B_ denote the variance ratio of SMA to GMA for sorghum and soybean, respectively.

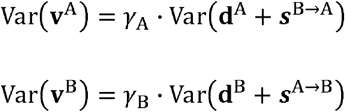
5. Finally, the residuals were sampled independently for each genotype in each pair from a normal distribution according to the specified heritability 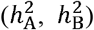 for each scenario. The variance of the residuals was calculated as follows:

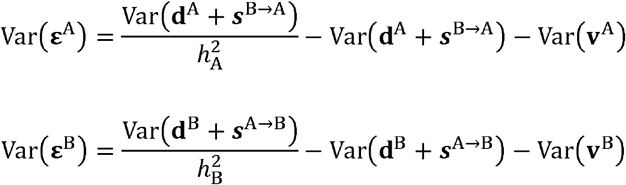

In this manner, pseudo-phenotypes were generated for each individual constituting the 5,214 genotype pairs in each scenario. For each of the 18 phenotype scenarios, to avoid bias arising from the positions and effects of the QTNs, the QTN positions and effects at step 1 were resampled 10 times, resulting in 10 phenotype sets for each scenario. To facilitate comparisons among crops and scenarios in subsequent analyses, the pseudo-phenotypes were scaled separately for each crop within each scenario × phenotype replicate to have a mean of 0 and a variance of 1. The scaled pseudo-phenotypes were used in all subsequent analyses.

### 2.4. Genomic prediction model

Five genomic prediction (GP) models were applied to the scaled pseudo-phenotypes and the prediction accuracy of each model was evaluated. The mathematical formulations for each model are presented below.

Model 1:

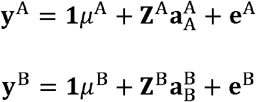

Model 2:

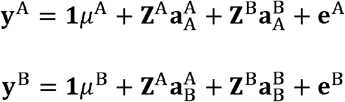

Model 3:

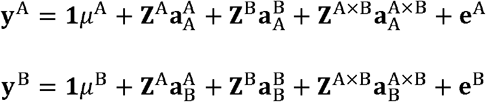

Here, **y**^A^ and **y**^B^ are 5214 × 1 vectors indicating the pseudo-phenotypes of crop A (sorghum) and crop B (soybean), respectively, generated for each intercropping pair. *μ*^A^ and *μ*^B^ are the overall means. **e**^A^ and **e**^B^ are errors following the distributions 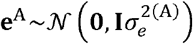 and 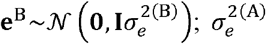 and 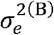 are the error variances. **Z**^A^ and **Z**^A^ are incidence matrices linking genotype effects to observations, whereas **Z**^AXB^ links pair-specific interaction effects (SMA) to each genotype pair, effectively assigning a unique interaction effect to each pair. 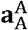 and 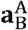 are 79 × 1 vectors representing the additive genetic effects of crop A, and are assumed to follow the distributions 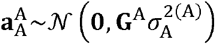 and 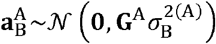.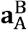 and 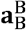 are 66 × 1 vectors representing the additive genetic effects of crop B, and are assumed to follow the distributions 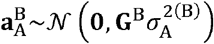 and 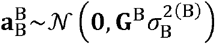. **G**^A^ and **G**^B^ are additive genomic relationship matrices for A and B, respectively, and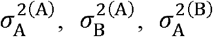 and 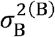 are the additive genetic variances. 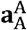 and 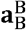 represent the additive genetic efforts of A and B on themselves, respectively (i.e., DGEs), whereas 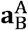 and 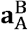 represent the additive genetic effects exerted by the paired genotype (i.e., IGEs). In addition. 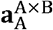 and 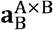 are 5214 × 1 vectors representing the SMA, and are assumed to follow the distributions 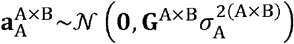 and 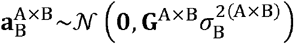. Here, **G** ^A×B^ is a 5214 × 5214 matrix given by the Kronecker product of **G**^A^ and **G**^B^ to represent pairwise genetic relationships between genotypes of the two species; 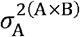 and 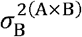 denote the variance components of the SMA.

Models 4 and 5 are multivariate extensions of Models 2 and 3, respectively. In both models, unstructured covariances between 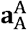 and 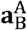, between 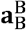 and 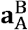, and between **e**^A^ and **e**^B^ were allowed. In Model 5, however, the covariance between 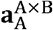 and 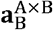 was set to zero.

Overall, Models 1 to 3 were fitted separately for each crop. Among these models, Model 1 considered only DGE, Model 2 considered both DGE and IGE, and Model 3 incorporated SMA in addition to DGE and IGE. Models 4 and 5 treated the phenotypes of sorghum and soybean as two traits and extended Models 2 and 3, respectively, into multivariate models that allowed genetic correlations between DGE and IGE. Models 1–5 were fitted using the R package ‘BGLR’ (version 1.1.4) (Pérez and de los Campos, 2014; Pérez-Rodríguez and de los Campos, 2022). For estimation of random effects, the reproducing kernel Hilbert space model was used. Models 1–3 were fitted using the ‘BGLR’ function, whereas Models 4 and 5 were fitted using the ‘Multitrait’ function. The Markov chain Monte Carlo was run for 10,000 iterations, with the first 2,000 samples discarded as burn-in and a sampling interval (thinning) of 20.

### 2.5. Assessment of genomic prediction accuracy

For each of the 18 scenarios and each of the 10 replicated phenotype sets, the accuracy of predictions with Models 1–5 was evaluated by cross-validation. In each cross-validation run, 520 of the 5,214 intercropping pairs were selected as the training set and the remaining 4,694 pairs were used as the test set. Model performance was assessed by calculating the Pearson correlation coefficient between the true phenotypic values, defined as the pseudo-phenotypes generated in Section 2.3, and the predicted phenotypic values in the test set. The use of 520 pairs for model training was intended to reflect the approximate capacity of a realistic field experiment. In addition, two training-set designs were considered (Figure S2). The first set, referred to as CV1, comprised all possible pairs between 26 sorghum accessions and 20 soybean accessions (520 pairs in total). The second set, referred to as CV2, comprised 520 pairs such that all genotypes were represented as evenly as possible across pairs: each sorghum accession appeared six or seven times, and each soybean accession appeared seven or eight times. CV1 represents a strategy based on dense evaluation of a restricted set of genotypes, whereas CV2 represents a strategy based on broad genotype coverage under limited evaluation capacity. This comparison allowed evaluation of which strategy is more effective for genomic prediction in intercropping systems.

For each scenario and each cross-validation training set, the selection of the 520 training pairs was repeated 10 times. The prediction accuracy was averaged over the resulting 100 runs, corresponding to 10 phenotype replicates and 10 training-set replicates.

### 2.6. Bayesian optimization

Figure 1 presents a schematic overview of the Bayesian optimization procedure used. For a given intercropping pair, the total genotypic values of crops A (sorghum) and B (soybean) in Model 3 are expressed as follows.

**Figure 1.**
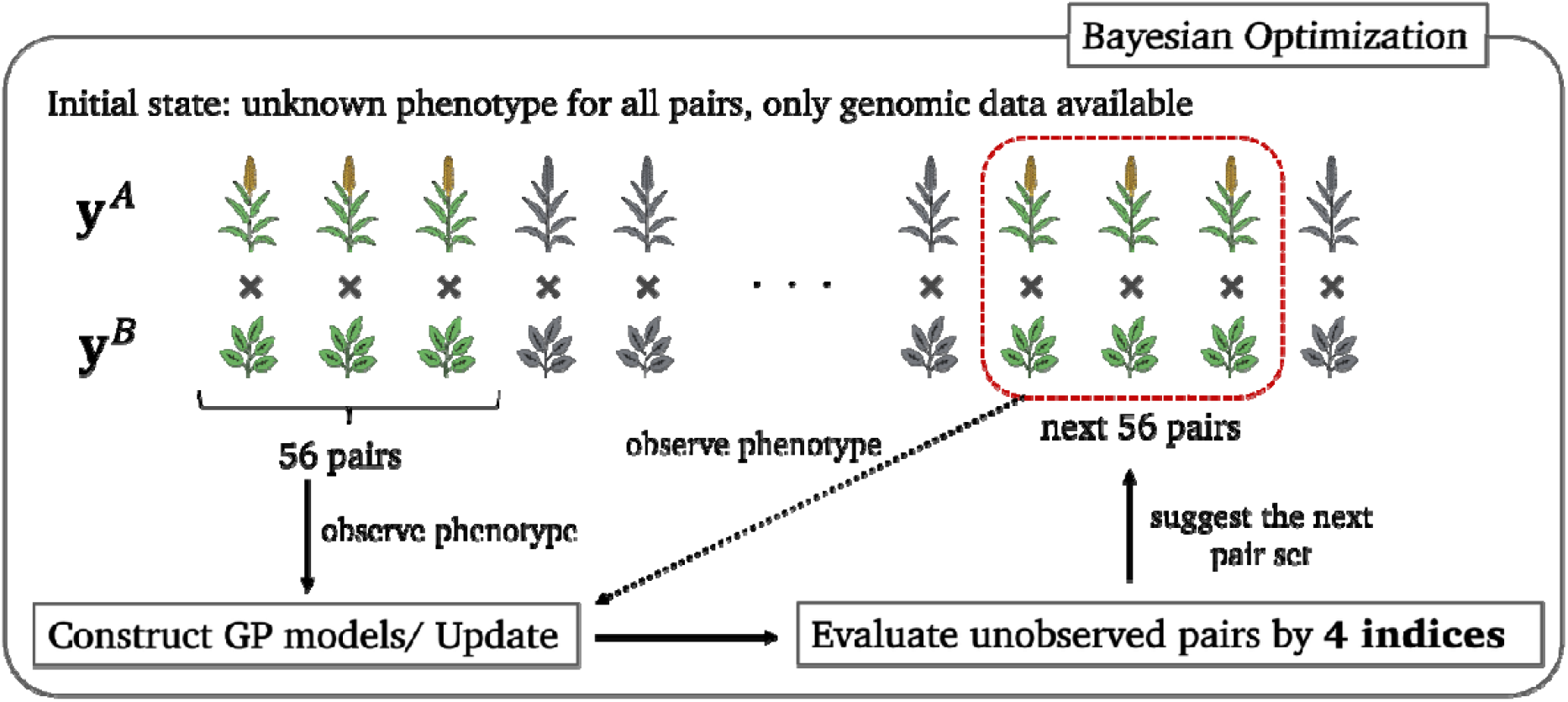
Schematic overview of the Bayesian optimization procedure used in this study.

where,, and are the th row of,, and, respectively.

The objective function used for Bayesian optimization was defined as the sum of the two crops’ total genotypic values, i.e.,. Prior to starting the optimization procedure, the phenotypic data were masked for all 5,214 genotype pairs formed between the 79 sorghum accessions and the 66 soybean accessions, while SNP data were available for all accessions of both crops. From these genotype pairs, 56 were randomly selected and the scaled pseudo-phenotypes (generated as described in Section 2.3) were selected. Using these 56 observations together with the corresponding SNP data, a genomic prediction model was constructed using Model 3. Based on the variance components estimated by Model 3, four indices described below were calculated for the remaining 5,158 unobserved pairs. The 56 pairs with the highest index values were selected and added to the genomic prediction model to update it. At each cycle, including the initial step, a fixed number of pairs (56) were evaluated, corresponding to a realistic per-cycle evaluation capacity in field experiments. This cycle was repeated until the optimal pair, defined as the pair with the highest total true genotypic value, was identified.

At optimization cycle *t*, given the observed data 𝒟*t* the posterior predictive distribution of the objective value for pair *k* is described as 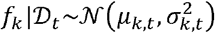, where *μ*_*k,t*_ and *σ*_*k,t*_ denote the posterior mean and posterior standard deviation, respectively. The derivation of *μ*_*k,t*_ and 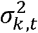 is provided in Supplementary file 2. The four acquisition functions described below were used to prioritize the remaining unobserved pairs and thereby guide the search for the optimal solution.

Posterior mean (Mu):*α*_M u_ (*k*) = *μ*_*k,t*_

Probability of Improvement (PI):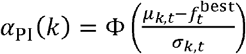

Expected Improvement (EI): 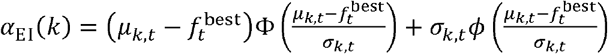

Upper Confidence Bound (UCB): *α*_UCB_(*k*) = *μ*_*k,t* +_ *k σ*_*k,t*_

Where 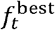 is the best objective value observed up to cycle *t*, and *k* > 0 is an exploration parameter in the UCB formula that determines the quantile of the posterior predictive distribution. In this study, we set *k* =1.64, corresponding to the 95th percentile of the posterior predictive distribution. Φ(·) and *ϕ* (·) are the cumulative distribution function and probability density function of the standard normal distribution, respectively. Among these indices, PI, EI, and UCB are acquisition functions specific to Bayesian optimization.

For each phenotype scenario, Bayesian optimization was performed on 10 phenotype replicates. For each replicate, 10 independent optimization runs were conducted using different randomly selected initial sets of 56 pairs, resulting in 100 runs per scenario. In addition, the initial random selection was constrained so that the optimal pair was not included.

### 2.7. Expected number of cycles under a random search

As a baseline, we derived the expected number of cycles required to identify the optimal pair under a random search. In each cycle, 56 pairs were sampled uniformly at random without replacement from the set of pairs that had not yet been evaluated. Let *T* denote the cycle at which the optimal pair is first identified. Because there is exactly one optimal pair among the 5,214 candidates, and each pair is equally likely to appear in any cycle under random sampling without replacement, the probability that it is identified at cycle *x* is constant for *x* =1, …, 93, namely,

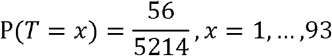

where 93 × 56 = 5208. For the final cycle, only the remaining six pairs are evaluated, and therefore

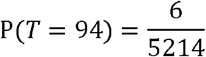

Thus, the expected number of cycles until the optimal pair is identified is given by

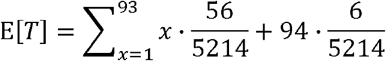

Because 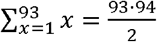, we obtained E[*T*] = 47.054.

Therefore, under a random search, the expected number of cycles required to identify the optimal pair is 47.054.

### 2.8. Optimization evaluation

For optimization of the intercropping pairs, we evaluated the optimization performance using four summary statistics: Spearman’s rank correlation coefficients, the top-56 overlap, relative gap from the maximum (RelGap), and minimum distance to the observed set (MinDist).

Spearman’s rank correlation coefficients were calculated, at each cycle within a single optimization run, between (i) the predicted posterior mean of the total genotypic values and (ii) the corresponding true values of the 56 pairs selected in that cycle. Here, a single optimization run denotes one of the 100 independent runs defined by phenotype replication × BO replication. The total genotypic value was defined as the sum of the genotypic values of the two individuals constituting an intercropping pair, and its predicted posterior mean was calculated as described in Section 2.6.

Top-56 overlap was defined as the number of pairs shared between the 56 pairs selected in a given cycle and the top 56 unobserved pairs ranked by true total genotypic value at that cycle.

Relative gap from the maximum (RelGap) was calculated as follows: 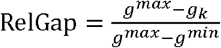 RelGap was calculated for each cycle within a single optimization run. Here, *k* denotes one of the 56 pairs selected in a given cycle, and *g*_*k*_ denotes the true total genotypic value of pair *k. g*^*max*^ and *g*^*min*^ respectively denote the maximum and minimum true total genotypic values among the 5,214 pairs in that phenotype replicate. This definition of RelGap enabled comparisons across phenotype replicates with different value ranges.

Minimum distance to the observed set (MinDist) was calculated as follows: *MinDist* = min_j ∈ 𝒪_ where 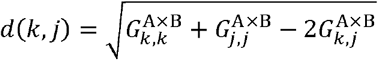. MinDist was calculated for each cycle within a single optimization run. Here, *k* denotes one of the 56 pairs selected in a given cycle, 𝒪 denotes the set of pairs observed before that cycle, and *j* denotes one pair in 𝒪.*G*^A × B^ is a 5,214 × 5,214 matrix given by the Kronecker product of the genomic relationship matrices for sorghum and soybean.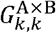 and 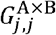are the diagonal elements corresponding to pairs *k* and *j* themselves, respectively, and 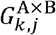 is the off-diagonal element representing the genetic relationship between pairs *k* and *j*. The distance *d* was defined so that genetically similar pairs had smaller values. Thus, MinDist quantifies how far each cycle explored new regions of the genetic space relative to the previously observed set.

## 3. Results

### 3.1. Accuracy of genomic prediction models

Figure 2 and Table S1 show the mean prediction accuracy achieved with the five genomic prediction models across the 18 phenotype scenarios and two training-set designs. Predictions were performed separately for sorghum and soybean; however, because no substantial differences were observed between the two crops, the results were averaged across the two crops. With respect to the training-set design, CV2 (in which all genotypes were represented as evenly as possible) showed higher prediction accuracy than CV1 (in which all pairs from a subset of genotypes were selected). Regarding the phenotype scenarios, the prediction accuracy was higher when heritability was high. Changes in the variance ratio of SMA to GMA had little effect on prediction accuracy overall; however, for Models 3 and 5, which account for SMA, the prediction accuracy increased slightly as the variance ratio of SMA became larger. When the genetic correlation between DGE and IGE was varied, all models showed higher prediction accuracy under strong positive or negative correlations (−0.7 or 0.7) than under a near-zero correlation (−0.1).

**Figure 2.**
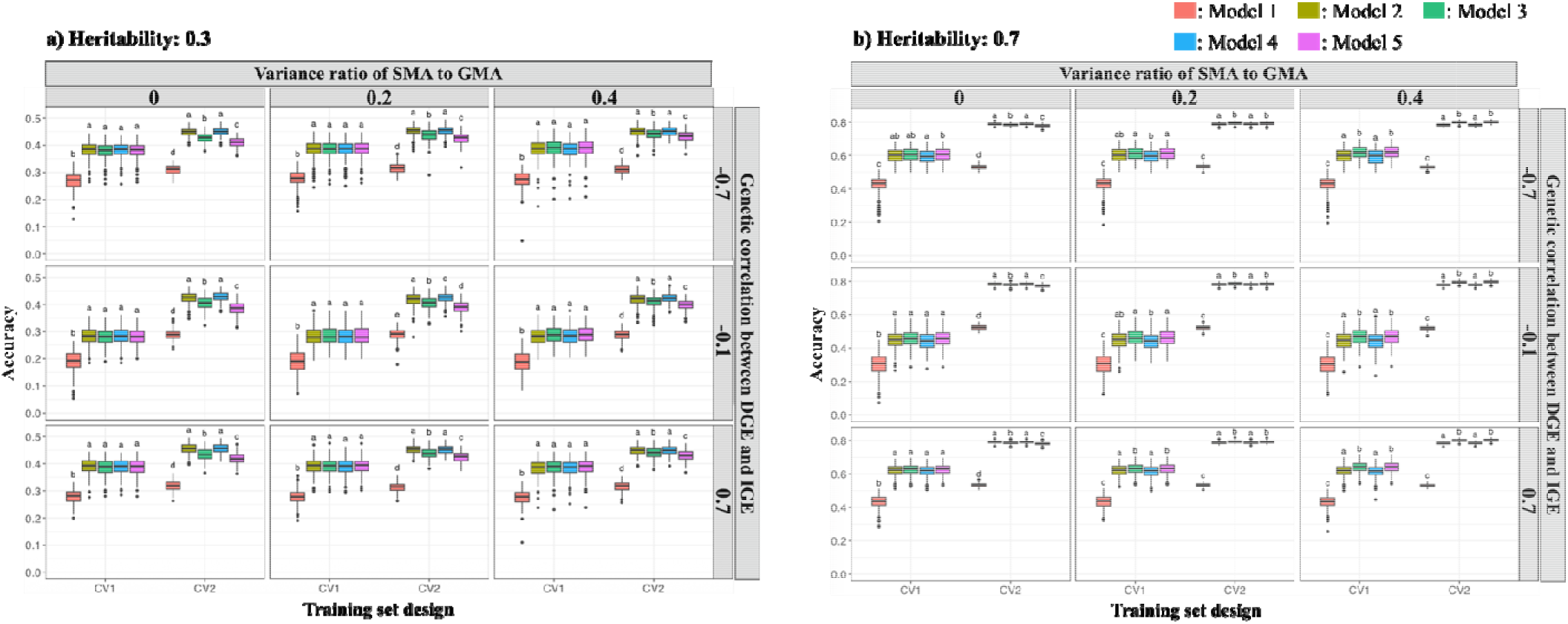
Mean prediction accuracy of five genomic prediction models across 18 phenotype scenarios and two training-set designs. Prediction accuracy was evaluated as the Pearson correlation between observed and predicted phenotype values. Panel (a) corresponds to *h*^2^ = 0.3 and panel (b) corresponds to *h*^2^ = 0.7. Within each panel, the results are presented according to the variance ratio of SMA to GMA (0, 0.2, and 0.4) and the genetic correlation between DGE and IGE (−0.7, −0.1, and 0.7). Different lowercase letters above boxes indicate significant differences among models within each scenario (*P* < 0.05; ANOVA followed by Tukey’s HSD test).

Among the genomic prediction models, Model 1, which considered only DGE, showed markedly lower prediction accuracy overall. Incorporating covariance between DGE and IGE did not lead to distinct improvements in prediction accuracy (Model 4 vs. Model 2, and Model 5 vs. Model 3). In contrast, the inclusion of SMA (Models 3 and 5 vs. Models 2 and 4) affected model performance depending on the scenario. As a result, Models 2 and 4 showed higher prediction accuracy in a limited number of scenarios, specifically when heritability was 0.3 under CV2 and when heritability was 0.7 under CV2 with no SMA contribution, whereas Models 3 and 5 consistently outperformed Models 2 and 4 when heritability was high, except when the variance ratio of SMA to GMA was 0.

### 3.2. Number of cycles required to identify the optimal pair in optimization

Figure 3 and Table S2 show, for each phenotype scenario, the mean number of cycles required for the optimization procedure to identify the pair with the highest total true genotypic value when different evaluation indices for intercropping pairs were used. The expected number of cycles required to identify the optimal pair under a random search was 47.054, whereas all criteria based on genomic prediction identified the optimal pair in substantially fewer cycles.

**Figure 3.**
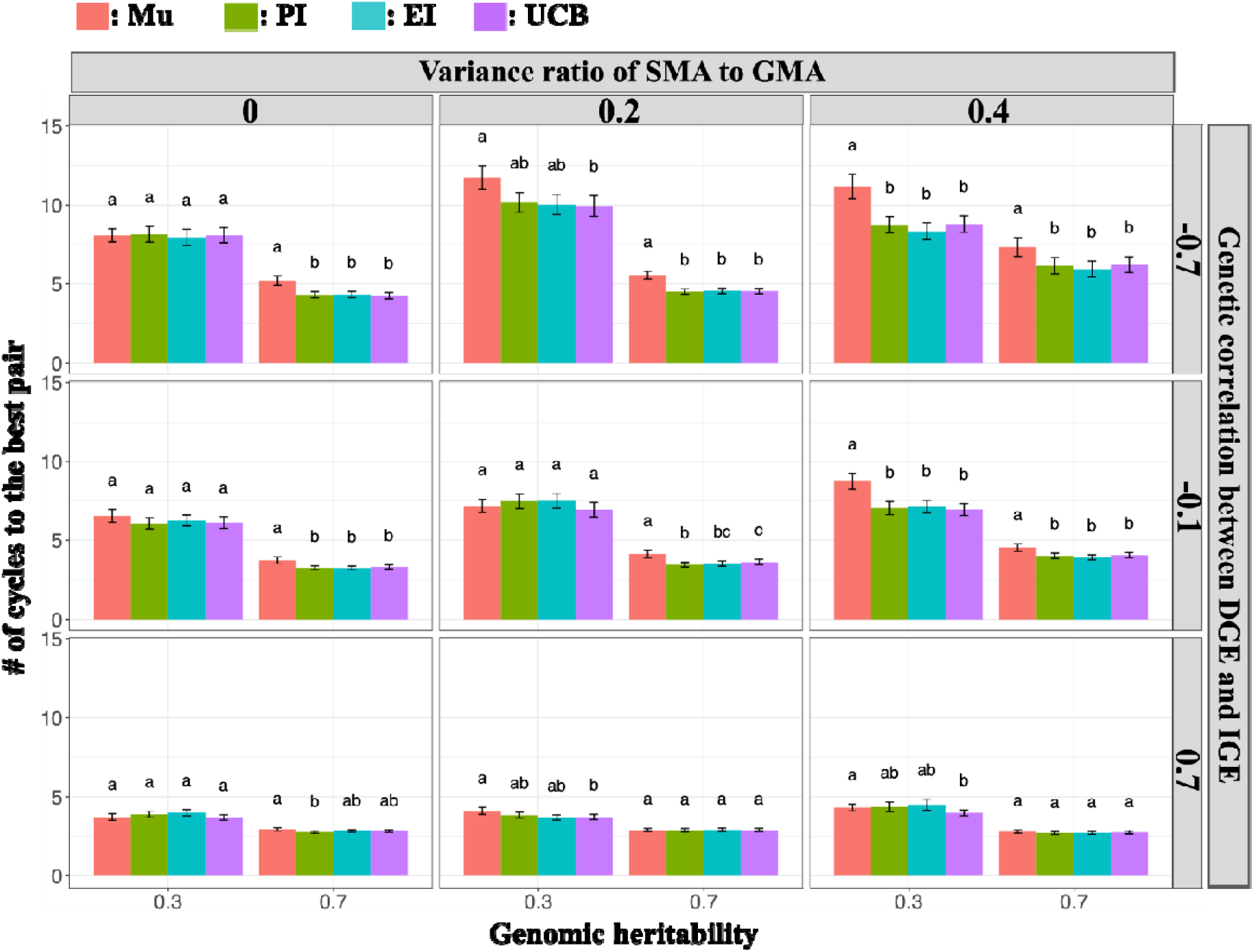
Mean number of cycles required to identify the optimal genotype pair using different evaluation indices for intercropping pairs (Mu, PI, EI, and UCB) across 18 phenotype scenarios. Lower values indicate more rapid identification of the optimal pair. Different lowercase letters above the bars indicate significant differences among evaluation indices within each scenario (*p* < 0.05; Friedman test followed by pairwise Wilcoxon tests).

Across all phenotype scenarios, the optimal pair was identified more rapidly when heritability was higher. When the variance ratio of SMA to GMA varied, the optimal pair was identified fastest in most scenarios when SMA was absent. With increase in the variance proportion of SMA to 0.2 and 0.4, the condition that identified the optimal pair most rapidly depended on the scenario; however, except in the cases where *h*^2^ = 0.3 and the genetic correlation between DGE and IGE was −0.1, and where *h*^2^ = 0.7 and the genetic correlation between DGE and IGE was 0.7, a smaller SMA variance generally resulted in faster identification of the optimal pair. When the genetic correlation between DGE and IGE varied, the number of cycles required to identify the optimal pair was shortest when the positive genetic correlation was strong, followed by when the correlation was close to zero, and longest under a strong negative correlation.

Comparison of the four evaluation indices for intercropping pairs showed that, except in certain scenarios, the three Bayesian optimization acquisition functions incorporating posterior variance in addition to the posterior mean of genomic prediction (PI, EI, and UCB) identified the optimal pair significantly faster than Mu, which used only the posterior mean. No significant difference from Mu was observed for the following scenarios: when SMA was absent and *h*^2^ = 0.3; when the SMA variance ratio was 0.2, the genetic correlation between DGE and IGE was −0.1, and *h*^2^ = 0.3; when the SMA variance ratio was 0.2, the genetic correlation was 0.7, and *h*^2^ = 0.7; and when the SMA variance ratio was 0.4, the genetic correlation was 0.7, and *h*^2^ = 0.7.

### 3.3. Summary statistics for optimization performance

Figures 4 and 5 show cycle-wise change in Spearman’s rank correlation coefficients and the top-56 overlap, as defined in Section 2.8. Regarding the Spearman’s rank correlation between predicted and true total genotypic values for the 56 pairs selected in each cycle, Mu and UCB showed lower Spearman’s rank correlation coefficients than PI and EI across all cycles in most phenotype scenarios. When the genetic correlation between DGE and IGE varied, a strong positive genetic correlation resulted in high Spearman’s rank correlation coefficients from the early cycles onward, which further increased over subsequent cycles. As the genetic correlation became more negative (i.e., decreased from to), the Spearman’s rank correlation coefficients decreased throughout the optimization process, and under a genetic correlation of the coefficients showed little improvement across cycles.

**Figure 4.**
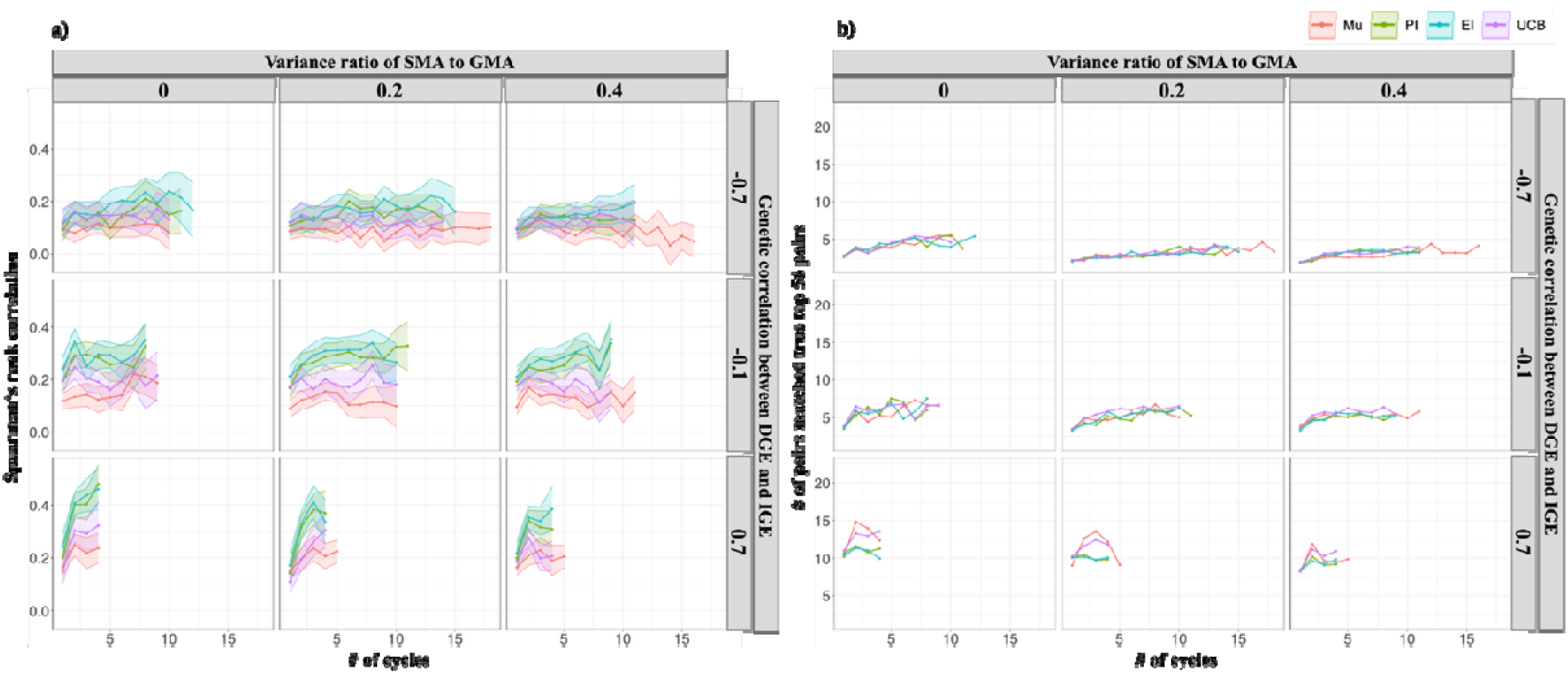
Changes in Spearman’s rank correlation coefficients and the top-56 overlap across optimization cycles for all phenotype scenarios with *h*^2^ = 0.3. (a) Spearman’s rank correlation coefficients and (b) the top-56 overlap. The *x*-axis origin is the cycle following the randomly selected initial pairs. Solid lines represent the mean across 100 optimization runs (10 phenotype replicates × 10 BO replicates). Shading in (a) represents the 95% confidence interval for the Spearman’s rank correlation coefficients. To avoid unstable summaries in later cycles, only cycles with at least 20 remaining runs are shown.

**Figure 5.**
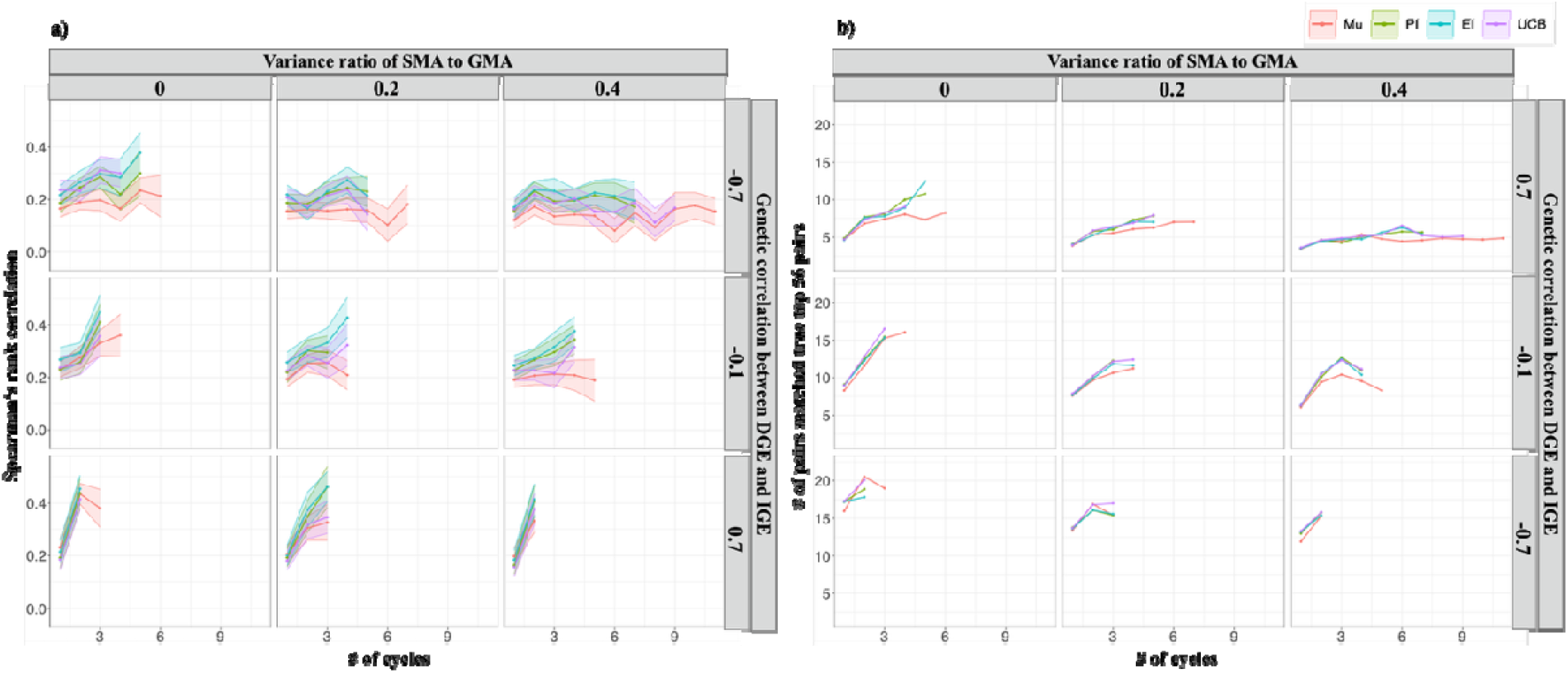
Changes in Spearman’s rank correlation coefficients and the top-56 overlap across optimization cycles for all phenotype scenarios with *h*^2^ = 0.7. (a) Spearman’s rank correlation coefficients and (b) the top-56 overlap. The *x*-axis origin is the cycle following the randomly selected initial pairs. Solid lines represent the mean across 100 optimization runs (10 phenotype replicates × 10 BO replicates). Shading in (a) represents the 95% confidence interval for the Spearman’s rank correlation coefficients. To avoid unstable summaries in later cycles, only cycles with at least 20 remaining runs are shown.

For the top-56 overlap, Mu and UCB showed greater overlap than PI and EI only in scenarios where *h*^2^ = 0.3 and the genetic correlation between DGE and IGE was 0.7; in the other scenarios, no distinct differences were observed among the four evaluation indices. Although overlap increased across cycles, overall, the top-56 overlap tended to increase in the order of genetic correlations of −0.7, −0.1, and 0.7.

Figure 6 shows the cycle-wise changes in RelGap and MinDist, as defined in Section 2.8, for the scenario with *h*^2^ = 0.3, variance ratio of SMA to GMA of 0.4, and genetic correlation between DGE and IGE of −0.7. Compared with Mu, the Bayesian optimization acquisition functions (PI, EI, and UCB) exhibited larger MinDist values throughout the optimization process. In addition, these acquisition functions explored broader regions of the genetic space throughout the optimization process, while gradually focusing on narrower regions in the later cycles. In contrast, Mu consistently explored regions genetically closer to the observed set.

**Figure 6.**
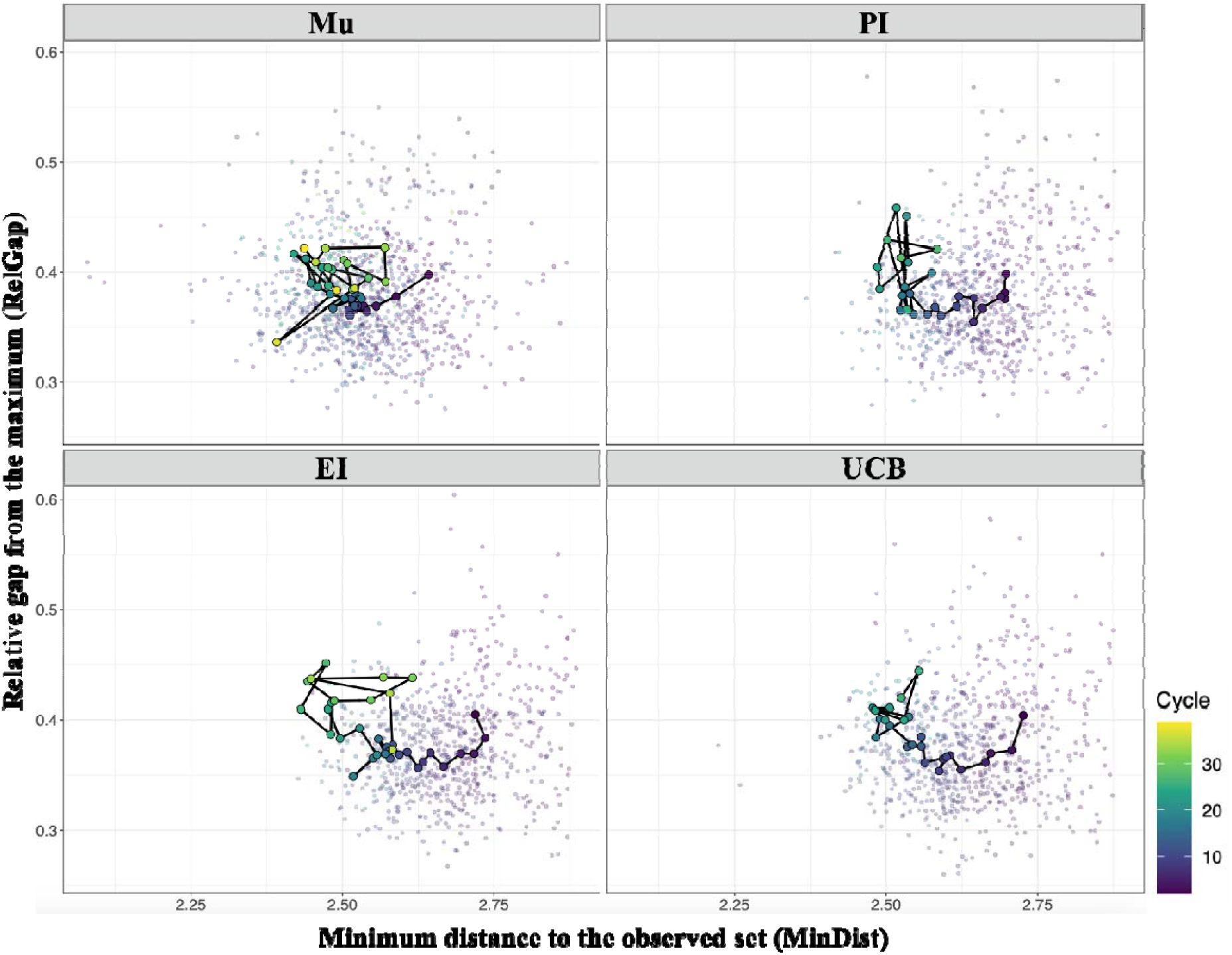
Cycle-wise changes in MinDist and RelGap for the scenario with *h*^2^ = 0.3, variance ratio of SMA to GMA = 0.4, and DGE–IGE genetic correlation =. The panels show the results for four evaluation indices (Mu, PI, EI, and UCB). Semi-transparent points represent the median MinDist and median RelGap for the 56 pairs selected in each cycle within each phenotype replicate × BO replicate combination. Black lines with colored-edge points represent cycle-wise medians across runs. The color indicates the cycle number.

## 4. Discussion

In this study, we evaluated the utility of a method that combines genomic prediction with Bayesian optimization for efficient identification of optimal intercropping pairs from existing genetic resources with known genomic information. To assess the accuracy of the genomic predictions, we compared different training-set designs and models. Increasing the number of genotypes included in the training set improved the prediction accuracy for untested pairs to a greater degree than evaluating all possible pairs within a limited subset of genotypes (Figure 2). This finding suggests that, even when the size of the training data is unchanged, the prediction accuracy is improved when the training set contains individuals that are more closely related to the target individuals, and that it is more important for the structure of the training data to adequately cover the genetic space of the prediction targets (Lorenz and Smith, 2015; Lee et al., 2017). However, although it may be feasible to evaluate all pairs among a limited number of genotypes, it may be difficult in practice to assemble a broad range of genotypes. Therefore, in future studies, identification of an appropriate balance between the number of genotypes included in the training set and the resulting prediction accuracy is required.

Among the five genomic prediction models assessed, Model 1, which considered only DGE, showed markedly lower prediction accuracy because it predicted each genotype based only on its own average genetic effect, regardless of the paired genotypes. Moreover, the multivariate models, Models 4 and 5, which incorporated a covariance structure between DGE and IGE, did not improve prediction accuracy relative to Models 2 and 3. Previous studies have suggested that multivariate models are particularly advantageous when a genetically correlated secondary trait is observed in the test set while the focal trait is missing (Arojju et al., 2020; Gill et al., 2021). In the present study, however, phenotypes were evaluated on a pairwise basis, and the phenotypes of both crops in a given pair were observed simultaneously. Therefore, the present assessment did not correspond to the situation in which only a genetically correlated secondary trait is available in the test set, which may explain the limited advantage of the multivariate models. The superiority of Model 3 over Model 2 under high-heritability scenarios suggests that the variance explained by SMA was sufficiently large relative to the residual variation to be captured by the model. Thus, when the genetic determination was strong, explicitly modeling SMA enabled the model to account for pair-specific genetic deviations beyond DGE and IGE, thereby improving the prediction accuracy. This interpretation is consistent with previous reports that models incorporating non-additive or interaction-related genetic components can outperform simpler additive models when such components contribute meaningfully to the underlying genetic architecture (Jiang and Reif, 2015; Morgante et al., 2018).

The use of genomic prediction and Bayesian optimization to identify the optimal intercropping pair was substantially more efficient than a random search, which is consistent with previous findings that these approaches can accelerate the identification of promising lines from pre-breeding populations (Tanaka and Iwata, 2018). Scenarios in which no difference was observed in the number of cycles required to identify the optimal pair between the posterior mean (Mu) and the three BO acquisition functions were those not considering SMA, or those with high heritability and a strong positive genetic correlation between DGE and IGE (Figure 3). In these cases, the optimization landscape defined by the objective function was either relatively simple because it did not include interaction effects, or the benefit of exploring unknown regions was limited, such that the predictive mean alone was sufficient. Indeed, Figures 4 and 5 show that, in scenarios with high heritability or a strong positive genetic correlation between DGE and IGE, the Spearman’s rank correlation coefficients between the predicted posterior mean and the true total genotypic values for the 56 pairs selected in each cycle increased more rapidly in the early cycles than in the other scenarios. A similar pattern was observed for the top-56 overlap, indicating that these scenarios enabled earlier concentration of the search on truly superior pairs. Previous research has suggested that, when differences in uncertainty among candidate points are small, the advantage of using Bayesian optimization acquisition functions that account for posterior variance becomes limited (Tsai et al., 2021). By contrast, scenarios in which the BO acquisition functions were advantageous were those in which SMA was present, making the objective function more complex through pair-specific interaction effects. This advantage was particularly pronounced when the genetic correlation between DGE and IGE was strongly negative; under such a scenario, even if a genotype has a positive effect on its own phenotype through DGE, it exerts an opposite effect on the paired genotype. Therefore, when the objective function is defined as the sum of the two crops, it becomes necessary to identify pairs in which the genetic effect on the individual itself and the genetic effect on its pair, acting in opposing directions, are appropriately balanced. Under such conditions, exploration of genetically unexplored regions becomes especially important. In addition, as shown in Figures 4 and 5, the Spearman’s rank correlation coefficients tended to be higher for PI and EI, which place greater emphasis on predictive uncertainty, than for Mu and UCB. This pattern may reflect improvements in the prediction accuracy of the genomic prediction model during the optimization process. Because PI and EI actively explore genetically distant regions with high uncertainty, the datasets used to update the prediction model likely captured broader genetic diversity. Such broader genetic coverage may have contributed to the improved model accuracy, resulting in greater agreement between the predicted posterior means and the true total genotypic values of the selected pairs. Therefore, PI and EI appear to facilitate exploration while simultaneously improving predictive performance, suggesting that these acquisition functions are well suited for identifying promising pairs throughout the optimization process.

In this study, the objective function for optimization was defined as the sum of the genotypic values of the two crops. However, depending on the intercropping context, it may sometimes be more appropriate to prioritize the phenotype of only one crop or to optimize the phenotypes of both crops in a more balanced manner. Therefore, depending on the breeding objective, extensions such as multi-objective Bayesian optimization may be warranted.

This study demonstrated the potential utility of genomic prediction for identifying optimal genotype pairs in interspecific intercropping systems. In addition, we evaluated a Bayesian optimization approach that incorporates the posterior mean of genomic prediction and the predictive variance. Simulation-based analyses showed that this approach was particularly advantageous under complex intercropping genetic architectures, such as when SMA contributed substantially or when the genetic correlation between DGE and IGE was strongly negative. Future studies should evaluate the practical effectiveness of this framework in combination with empirical studies and clarify challenges that arise in field-based intercropping experiments.

## Acknowledgements

We gratefully acknowledge Dr. Hideki Takanashi and Dr. Hiromi Kajiya-Kanegae for providing the genomic data for 237 sorghum germplasm accessions, and Dr. Akito Kaga and Dr. Hiromi Kajiya-Kanegae for providing the genomic data for 198 soybean germplasm accessions. We thank Robert McKenzie, PhD, from Edanz (https://jp.edanz.com/ac) for editing the English text of a draft of this manuscript.

## Funding statement

This work was supported by the Japan Science and Technology Agency (JST) Strategic Basic Research Programs, Advanced Technologies for Carbon-Neutral (ALCA-Next) [Grant Number JPMJAN23D1]. The funder had no role in the study design, data collection and analysis, the decision to publish, or preparation of the manuscript.

## Author contributions

All authors contributed to the conception and design of this study. S.K.: Conceptualization, Data curation, Methodology, Formal analysis, Software, Visualization, Writing – original draft. H.I.: Conceptualization, Methodology, Funding acquisition, Supervision, Project administration, Writing – review & editing.

## Data availability

The datasets and R code used in this study are available from the GitHub repository ‘Sei-Kinoshita/ICBO’ (https://github.com/Sei-Kinoshita/ICBO).

## Declaration of competing interests

The authors have no relevant financial or nonfinancial interests to disclose.

## Notes

### Competing Interest Statement

The authors have declared no competing interest.

## References

Abou Khater, L., Maalouf, F., Balech, R., He, Y., Zong, X., Rubiales, D., Kumar, S., 2024. Improvement of cool-season food legumes for adaptation to intercropping systems: Breeding faba bean for intercropping with durum wheat as a case study. Front. Plant Sci., 15, 1368509. 10.3389/fpls.2024.1368509

Akchaya, K., Parasuraman, P., Pandian, K., Vijayakumar, S., Thirukumaran, K., Mustaffa, M.R.A.F., Rajpoot, S.K., Choudhary, A.K., 2025. Boosting resource use efficiency, soil fertility, food security, ecosystem services, and climate resilience with legume intercropping: A review. Front. Sustain. Food Syst., 9, 1527256. 10.3389/fsufs.2025.1527256

Annicchiarico, P., Nazzicari, N., Notario, T., Monterrubio Martin, C., Romani, M., Ferrari, B., Pecetti, L., 2021. Pea breeding for intercropping with cereals: Variation for competitive ability and associated traits, and assessment of phenotypic and genomic selection strategies. Front. Plant Sci., 12, 731949. 10.3389/fpls.2021.731949

Arojju, S.K., Cao, M., Trolove, M., Barrett, B.A., Inch, C., Eady, C., Stewart, A., Faville, M.J., 2020. Multi-trait genomic prediction improves predictive ability for dry matter yield and water-soluble carbohydrates in perennial ryegrass. Front. Plant Sci., 11, 1197. 10.3389/fpls.2020.01197

Bančič, J., Werner, C.R., Gaynor, R.C., Gorjanc, G., Odeny, D.A., Ojulong, H.F., Dawson, I.K., Hoad, S.P., Hickey, J.M., 2021. Modeling illustrates that genomic selection provides new opportunities for intercrop breeding. Front. Plant Sci., 12, 605172. 10.3389/fpls.2021.605172

Bijma, P., 2014. The quantitative genetics of indirect genetic effects: A selective review of modelling issues. Heredity, 112, 61–69. 10.1038/hdy.2013.15

Bourke, P.M., Evers, J.B., Bijma, P., van Apeldoorn, D.F., Smulders, M.J.M., Kuyper, T.W., Mommer, L., Bonnema, G., 2021. Breeding beyond monoculture: Putting the “intercrop” into crops. Front. Plant Sci., 12, 734167. 10.3389/fpls.2021.734167

Diot, J., Iwata, H., 2023. Bayesian optimisation for breeding schemes. Front. Plant Sci., 13, 1050198. 10.3389/fpls.2022.1050198

Dubey, R., Zustovi, R., Landschoot, S., Dewitte, K., Verlinden, G., Haesaert, G., Maenhout, S., 2024. Harnessing monocrop breeding strategies for intercrops. Front. Plant Sci., 15, 1394413. 10.3389/fpls.2024.1394413

Firmat, C., Litrico, I., 2022. Linking quantitative genetics with community-level performance: Are there operational models for plant breeding? Front. Plant Sci., 13, 733996. 10.3389/fpls.2022.733996

Forst, E., Enjalbert, J., Allard, V., Ambroise, C., Krissaane, I., Mary-Huard, T., Robin, S., Goldringer, I., 2019. A generalized statistical framework to assess mixing ability from incomplete mixing designs using binary or higher order variety mixtures and application to wheat. Field Crops Res., 242, 107571. 10.1016/j.fcr.2019.107571

Gill, H.S., Halder, J., Zhang, J., Brar, N.K., Rai, T.S., Hall, C., Bernardo, A., St Amand, P., Bai, G., Olson, E., Ali, S.A., Turnipseed, B., Sehgal, S.K., 2021. Multi-trait multi-environment genomic prediction of agronomic traits in advanced breeding lines of winter wheat. Front. Plant Sci., 12, 709545. 10.3389/fpls.2021.709545

Haug, B., Messmer, M.M., Enjalbert, J., Goldringer, I., Forst, E., Flutre, T., Mary-Huard, T., Hohmann, P., 2021. Advances in breeding for mixed cropping—Incomplete factorials and the producer/associate concept. Front. Plant Sci., 11, 620400. 10.3389/fpls.2020.620400

Haug, B., Messmer, M.M., Enjalbert, J., Goldringer, I., Flutre, T., Mary-Huard, T., Hohmann, P., 2023. New insights towards breeding for mixed cropping of spring pea and barley to increase yield and yield stability. Field Crops Res., 297, 108923. 10.1016/j.fcr.2023.108923

Harwood, J., 2024. The forgotten history of intercropping. Plants, People, Planet 6(5), 1121–1128. 10.1002/ppp3.10502

Hohmann, P., Kussmann, S., Schöb, C., Rubiales, D., Annicchiarico, P., 2026. Rethinking variety testing to recognise mixing ability: Bridging breeding, policy, and practice in diversified agriculture. Front. Plant Sci., 17, 1770790. 10.3389/fpls.2026.1770790

Jiang, Y., Reif, J.C., 2015. Modeling epistasis in genomic selection. Genetics 201(2), 759–768. 10.1534/genetics.115.177907

Jing, H., Liu, Y., Hou, J., 2025. Impacts of agricultural intensification on biodiversity: Habitat loss, agrochemical use, water depletion, and soil degradation. J. Environ. Manage. 395, 128036. 10.1016/j.jenvman.2025.128036

Kaga, A., Shimizu, T., Watanabe, S., Tsubokura, Y., Katayose, Y., Harada, K., Vaughan, D.A., Tomooka, N., 2012. Evaluation of soybean germplasm conserved in NIAS genebank and development of mini core collections. Breed. Sci. 61(5), 566–592. 10.1270/jsbbs.61.566

Kajiya-Kanegae, H., Nagasaki, H., Kaga, A., Hirano, K., Ogiso-Tanaka, E., Matsuoka, M., Ishimori, M., Ishimoto, M., Hashiguchi, M., Tanaka, H., Akashi, R., Isobe, S., Iwata, H., 2021. Whole-genome sequence diversity and association analysis of 198 soybean accessions in mini-core collections. DNA Res. 28(1), dsaa032. 10.1093/dnares/dsaa032

Lee, K., Kim, M.-S., Lee, J.S., Bae, D.N., Jeong, N., Yang, K., Lee, J.-D., Park, J.-H., Moon, J.-K., Jeong, S.-C., 2020. Chromosomal features revealed by comparison of genetic maps of *Glycine max* and *Glycine soja*. Genomics 112(2), 1481–1489. 10.1016/j.ygeno.2019.08.019

Lee, S.H., Clark, S., van der Werf, J.H.J., 2017. Estimation of genomic prediction accuracy from reference populations with varying degrees of relationship. PLoS ONE 12(12), e0189775. 10.1371/journal.pone.0189775

Li, C., Stomph, T.-J., Makowski, D., Li, H., Zhang, C., Zhang, F., van der Werf, W., 2023. The productive performance of intercropping. Proc. Natl. Acad. Sci. U. S. A. 120(2), e2201886120. 10.1073/pnas.2201886120

Lorenz, A.J., Smith, K.P., 2015. Adding genetically distant individuals to training populations reduces genomic prediction accuracy in barley. Crop Sci. 55(6), 2657–2667. 10.2135/cropsci2014.12.0827

Mace, E.S., Rami, J.-F., Bouchet, S., Klein, P.E., Klein, R.R., Kilian, A., Wenzl, P., Xia, L., Halloran, K., Jordan, D.R., 2009. A consensus genetic map of sorghum that integrates multiple component maps and high-throughput Diversity Array Technology (DArT) markers. BMC Plant Biol. 9, 13. 10.1186/1471-2229-9-13

Moore, V.M., Schlautman, B., Fei, S.-Z., Roberts, L.M., Wolfe, M., Ryan, M.R., Wells, S., Lorenz, A.J., 2022. Plant breeding for intercropping in temperate field crop systems: A review. Front. Plant Sci. 13, 843065. 10.3389/fpls.2022.843065

Morgante, F., Huang, W., Maltecca, C., Mackay, T.F.C., 2018. Effect of genetic architecture on the prediction accuracy of quantitative traits in samples of unrelated individuals. Heredity 120, 500–514. 10.1038/s41437-017-0043-0

Neyhart, J.L., Lorenz, A.J., Smith, K.P., 2019. Multi-trait improvement by predicting genetic correlations in breeding crosses. G3: Genes, Genomes, Genet. 9(10), 3153–3165. 10.1534/g3.119.400406

Paterson, A.H., Bowers, J.E., Bruggmann, R., Dubchak, I., Grimwood, J., Gundlach, H., Haberer, G., Hellsten, U., Mitros, T., Poliakov, A., Schmutz, J., Spannagl, M., Tang, H., Wang, X., Wicker, T., Bharti, A.K., Chapman, J., Feltus, F.A., Gowik, U., Grigoriev, I.V., Lyons, E., Maher, C.A., Martis, M., Narechania, A., Otillar, R.P., Penning, B.W., Salamov, A.A., Wang, Y., Zhang, L., Carpita, N.C., Freeling, M., Gingle, A.R., Hash, C.T., Keller, B., Klein, P., Kresovich, S., McCann, M.C., Ming, R., Peterson, D.G., Mehboob-ur-Rahman Ware, D., Westhoff, P., Mayer, K.F.X., Messing, J. Rokhsar, D.S., 2009. The Sorghum bicolor genome and the diversification of grasses. Nature 457, 551–556. 10.1038/nature07723

Pérez, P., de los Campos, G., 2014. Genome-wide regression and prediction with the BGLR statistical package. Genetics 198(2), 483–495. 10.1534/genetics.114.164442

Pérez-Rodríguez, P., de los Campos, G., 2022. Multitrait Bayesian shrinkage and variable selection models with the BGLR-R package. Genetics 222(1), iyac112. 10.1093/genetics/iyac112

Siska, M., Pajak, E., Rosenthal, K., del Rio Chanona, A., von Lieres, E., Helleckes, L.M., 2026. A guide to Bayesian optimization in bioprocess engineering. Biotechnol. Bioeng. 123(4), 805–830. 10.1002/bit.70129

Takanashi, H., Kajiya-Kanegae, H., Nishimura, A., Yamada, J., Ishimori, M., Kobayashi, M., Yano, K., Iwata, H., Tsutsumi, N., Sakamoto, W., 2022. *DOMINANT AWN INHIBITOR* encodes the ALOG protein originating from gene duplication and inhibits awn elongation by suppressing cell proliferation and elongation in sorghum. Plant Cell Physiol. 63(7), 901–918. 10.1093/pcp/pcac057

Tanaka, R., Iwata, H., 2018. Bayesian optimization for genomic selection: A method for discovering the best genotype among a large number of candidates. Theor. Appl. Genet. 131, 93–105. 10.1007/s00122-017-2988-z

Toker, P., Canci, H., Turhan, I., Isci, A., Scherzinger, M., Kordrostami, M., Yol, E., 2024. The advantages of intercropping to improve productivity in food and forage production – a review. Plant Prod. Sci. 27(3), 155–169. 10.1080/1343943X.2024.2372878

Tsai, S.-F., Shen, C.-C., Liao, C.-T., 2021. Bayesian optimization approaches for identifying the best genotype from a candidate population. J. Agric. Biol. Environ. Stat. 26, 519–537. 10.1007/s13253-021-00454-2

Tu, H.-N., Liao, C.-T., 2026. A modified Bayesian optimization approach for determining a training set to identify the best genotypes from a candidate population in genomic selection. J. Agric. Biol. Environ. Stat. 31, 1–22. 10.1007/s13253-024-00632-y

