## Supplemental File 1 for "Efficient Optimization of Genotype Pairs for Intercropping using Genomic Prediction and Bayesian Optimization"

**Supplementary File**

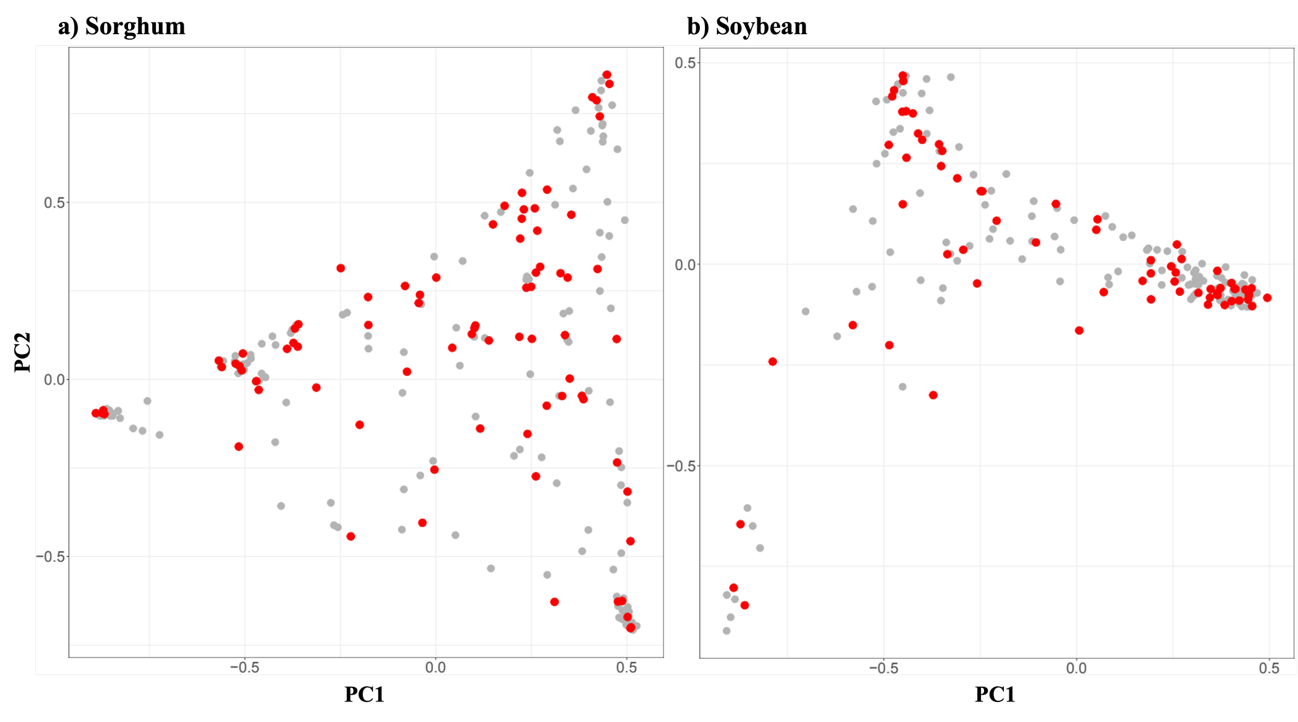

**Figure S1**. PCA of the 79 sorghum accessions and 66 soybean accessions selected by k-medoids. Principal component analysis was performed based on the eigendecomposition of the genomic relationship matrix for 237 sorghum accessions and 198 soybean accessions, and the results were plotted on PC1 and PC2. Panel (a) shows sorghum, and panel (b) shows soybean. Gray points indicate accessions not selected by k-medoids, whereas red points indicate accessions selected by k-medoids.

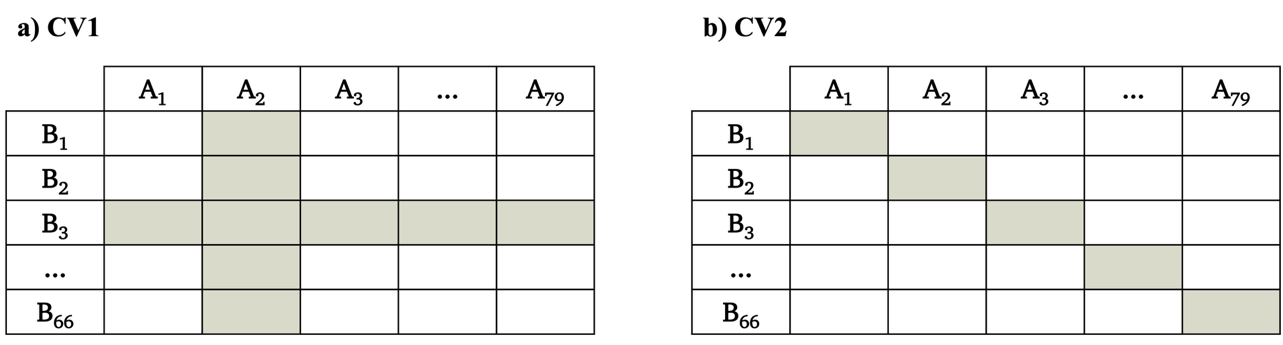

**Figure S2**. Schematic illustration of the two training-set designs. (a) CV1, in which all pairs of a subset of genotypes were selected as the training set; and (b) CV2, in which pairs were selected so that all genotypes were represented as evenly as possible and at least a certain number of times. A denotes sorghum and B denotes soybean. Colored cells indicate the pairs selected as the training set.

**TableS1.** Mean prediction accuracy of the five genomic prediction models for each phenotype scenario and training set design. Within each cell, the mean prediction accuracies of Models 1–5 are listed in order from top to bottom.

| **h^2^=0.3** | | | **h^2^=0.7** | | |  | | |
| --- | --- | --- | --- | --- | --- | --- | --- | --- |
| **Variance Ratio of SMA to GMA** | | | | | |  |  |  |
| **0** | **0.2** | **0.4** | **0** | **0.2** | **0.4** |  |  |  |
| 0.27  0.38  0.38  0.38  0.38 | 0.28  0.38  0.38  0.38  0.39 | 0.27  0.38  0.39  0.38  0.39 | 0.42  0.60  0.60  0.60  0.60 | 0.43  0.60  0.61  0.60  0.61 | 0.42  0.59  0.62  0.60  0.61 | **CV1** | **-0.7** | **Genetic Correlation between DGE and IGE** |
| 0.31  0.45  0.43  0.45  0.41 | 0.32  0.45  0.44  0.45  0.43 | 0.31  0.45  0.44  0.45  0.43 | 0.53  0.79  0.79  0.79  0.78 | 0.53  0.79  0.79  0.79  0.79 | 0.53  0.78  0.80  0.78  0.80 | **CV2** |  |  |
| 0.19  0.28  0.28  0.28  0.28 | 0.19  0.28  0.28  0.28  0.28 | 0.19  0.28  0.29  0.28  0.29 | 0.30  0.45  0.46  0.44  0.46 | 0.30  0.45  0.46  0.44  0.46 | 0.30  0.44  0.47  0.44  0.47 | **CV1** | **-0.1** |  |
| 0.29  0.43  0.40  0.43  0.39 | 0.29  0.42  0.40  0.42  0.39 | 0.29  0.42  0.41  0.42  0.40 | 0.52  0.78  0.78  0.78  0.77 | 0.52  0.78  0.79  0.78  0.78 | 0.52  0.78  0.80  0.78  0.80 | **CV2** |  |  |
| 0.28  0.39  0.39  0.39  0.39 | 0.28  0.39  0.39  0.39  0.39 | 0.27  0.38  0.39  0.38  0.39 | 0.43  0.62  0.63  0.62  0.63 | 0.43  0.62  0.63  0.62  0.63 | 0.43  0.62  0.64  0.62  0.64 | **CV1** | **0.7** |  |
| 0.32  0.46  0.44  0.46  0.42 | 0.31  0.45  0.44  0.45  0.43 | 0.32  0.45  0.44  0.45  0.43 | 0.53  0.80  0.79  0.79  0.78 | 0.53  0.79  0.80  0.79  0.79 | 0.53  0.78  0.80  0.78  0.80 | **CV2** |  |  |

**TableS2.** Mean number of cycles required to identify the optimal intercropping pair using different evaluation indices (Mu, PI, EI, and UCB) across phenotypic scenarios..

| **h^2^=0.3** | | | **h^2^=0.7** | | |  | | |
| --- | --- | --- | --- | --- | --- | --- | --- | --- |
| **Variance Ratio of SMA to GMA** | | | | | |  |  |  |
| **0** | **0.2** | **0.4** | **0** | **0.2** | **0.4** |  |  |  |
| 8.08 | 11.73 | 11.18 | 5.21 | 5.56 | 7.34 | **Mu** | **-0.7** | **Genetic Correlation between DGE and IGE** |
| 8.15 | 10.16 | 8.75 | 4.32 | 4.52 | 6.16 | **PI** |  |  |
| 7.95 | 10.03 | 8.34 | 4.33 | 4.56 | 5.94 | **EI** |  |  |
| 8.09 | 9.94 | 8.79 | 4.26 | 4.55 | 6.23 | **UCB** |  |  |
| 6.55 | 7.19 | 8.75 | 3.74 | 4.14 | 4.54 | **Mu** | **-0.1** |  |
| 6.07 | 7.48 | 7.04 | 3.26 | 3.46 | 4.02 | **PI** |  |  |
| 6.25 | 7.50 | 7.13 | 3.25 | 3.53 | 3.92 | **EI** |  |  |
| 6.12 | 6.94 | 6.95 | 3.32 | 3.63 | 4.06 | **UCB** |  |  |
| 3.72 | 4.11 | 4.32 | 2.95 | 2.90 | 2.80 | **Mu** | **0.7** |  |
| 3.89 | 3.84 | 4.36 | 2.77 | 2.89 | 2.71 | **PI** |  |  |
| 3.99 | 3.69 | 4.50 | 2.84 | 2.91 | 2.72 | **EI** |  |  |
| 3.69 | 3.72 | 3.98 | 2.83 | 2.90 | 2.75 | **UCB** |  |  |
