## Supplemental File 2 for "Efficient Optimization of Genotype Pairs for Intercropping using Genomic Prediction and Bayesian Optimization"

**Supplementary File 2**

**Derivation of posterior predictive mean and variance of total genotypic value under Model 3**

To calculate the 4 indices of evaluating intercropping pairs, we derived the posterior predictive mean and variance of the objective value for each unobserved pair under Model 3.

Let $\mathcal{O}$ denote the set of observed pairs, where $n_{\mathrm{obs}}$ is the number of observed pairs. Under Model 3, the phenotypic values of crop A and B in the observed pairs are modeled as

$$\mathbf{y}_{\mathcal{O}}^{A}=\boldsymbol{1}\mu^{A}+\mathbf{Z}_{\mathcal{O}}^{A}\mathbf{a}_{A}^{A}+\mathbf{Z}_{\mathcal{O}}^{B}\mathbf{a}_{A}^{B}+\mathbf{Z}_{\mathcal{O}}^{A\times B}\mathbf{a}_{A}^{A\times B}+\mathbf{e}_{\mathcal{O}}^{A}$$

$$\mathbf{y}_{\mathcal{O}}^{B}=\boldsymbol{1}\mu^{B}+\mathbf{Z}_{\mathcal{O}}^{A}\mathbf{a}_{B}^{A}+\mathbf{Z}_{\mathcal{O}}^{B}\mathbf{a}_{B}^{B}+\mathbf{Z}_{\mathcal{O}}^{A\times B}\mathbf{a}_{B}^{A\times B}+\mathbf{e}_{\mathcal{O}}^{B}$$

The posterior predictive distribution was derived separately for each crop using the corresponding Model 3, and the objective value was defined as the sum of the two crops’ total genotypic values. Therefore, below we describe the derivation for a general univariate Model 3. The same derivation applies to crop A and crop B. For a given crop,

$$\mathbf{y}_{\mathcal{O}}=\boldsymbol{1}\mu+\mathbf{u}_{\mathcal{O}}^{A}+\mathbf{u}_{\mathcal{O}}^{B}+\mathbf{u}_{\mathcal{O}}^{A\times B}+\mathbf{e}_{\mathcal{O}}$$

where $\mathbf{u}^{A}\boldsymbol{\sim}\mathcal{N}\left( \boldsymbol{0},\mathbf{K}^{A}\sigma^{2\left( A \right)} \right)$, $\mathbf{u}^{B}\boldsymbol{\sim}\mathcal{N}\left( \boldsymbol{0},\mathbf{K}^{B}\sigma^{2\left( B \right)} \right)$, $\mathbf{u}^{A\times B}\boldsymbol{\sim}\mathcal{N}\left( \boldsymbol{0},\mathbf{K}^{A\times B}\sigma^{2\left( A\times B \right)} \right)$, $\mathbf{e}\boldsymbol{\sim}\mathcal{N}\left( \boldsymbol{0},\mathbf{I}\sigma_{e}^{2} \right)$. Here, $\mathbf{K}^{A}=\mathbf{Z}^{A}\mathbf{G}^{A}\mathbf{Z}^{A^{T}}$, $\mathbf{K}^{B}=\mathbf{Z}^{B}\mathbf{G}^{B}\mathbf{Z}^{B^{T}}$, $\mathbf{K}^{A\times B}=\mathbf{Z}^{A\times B}\mathbf{G}^{A\times B}\mathbf{Z}^{{A\times B}^{T}}$ are the kernel matrices; $\sigma^{2\left( A \right)}$, $\sigma^{2\left( B \right)}$, $\sigma^{2\left( A\times B \right)}$, and $\sigma_{e}^{2}$ are the variance components estimated from Model 3.

Let $\mathbf{r}=\mathbf{y}_{\mathcal{O}}-\boldsymbol{1}\mu$. Then, the marginal distribution of the observed phenotypic values is $\boldsymbol{r\sim}\mathcal{N}\left( \boldsymbol{0},\mathbf{C} \right)$, where $\mathbf{C=}\mathbf{K}_{\mathcal{OO}}^{A}\sigma^{2\left( A \right)}+\mathbf{K}_{\mathcal{OO}}^{B}\sigma^{2\left( B \right)}+\mathbf{K}_{\mathcal{OO}}^{A\times B}\sigma^{2\left( A\times B \right)}+\mathbf{I}_{n_{\mathrm{obs}}}\sigma_{e}^{2}$. Here, $\mathbf{K}_{\mathcal{OO}}^{A}$, $\mathbf{K}_{\mathcal{OO}}^{B}$, and $\mathbf{K}_{\mathcal{OO}}^{A\times B}$ are the submatrices corresponding to observed pairs.

For an unobserved pair $k$, let $\mathbf{k}_{k\mathcal{O}}^{A}$**,** $\mathbf{k}_{k\mathcal{O}}^{B}$, and $\mathbf{k}_{k\mathcal{O}}^{A\times B}$ denote the $1\times n_{\mathrm{obs}}$ kernel vectors between pair $k$ and the observed pairs. Let the total genotypic value for pair $k$ be $g_{k}=u_{k}^{A}+u_{k}^{B}+u_{k}^{A\times B}$. Here, when the derivation is applied to crop A, this quantity corresponds to $g_{k}^{A}$ in the main text; when it is applied to crop B, it corresponds to $g_{k}^{B}$. Thus, $g_{k}$ here represents a crop-specific total genotypic value, whereas the Bayesian optimization objective is the sum of the two crop-specific values, $f_{k}=g_{k}^{A}\boldsymbol{+}g_{k}^{B}$. Then the variance of $g_{k}$ and the covariance between $g_{k}$ and $\mathbf{r}$ can be expressed as

$$\mathrm{Var}\left( g_{k} \right)=\mathbf{K}_{kk}^{A}\sigma^{2\left( A \right)}+\mathbf{K}_{kk}^{B}\sigma^{2\left( B \right)}+\mathbf{K}_{kk}^{A\times B}\sigma^{2\left( A\times B \right)}$$

$$\mathrm{Cov}\left( g_{k},\mathbf{r} \right)=\mathbf{k}_{k\mathcal{O}}^{A}\sigma^{2\left( A \right)}+\mathbf{k}_{k\mathcal{O}}^{B}\sigma^{2\left( B \right)}+\mathbf{k}_{k\mathcal{O}}^{A\times B}\sigma^{2\left( A\times B \right)}$$

Here, $\mathbf{K}_{kk}^{A}$, $\mathbf{K}_{kk}^{B}$, and $\mathbf{K}_{kk}^{A\times B}$ denote the diagonal elements of the three kernel matrices for pair $k$.

Then, the joint distribution of $g_{k}$ and $\mathbf{r}$ is

$$\left( \begin{aligned} g_{k} \\ \mathbf{r} \end{aligned} \right)\sim\mathcal{N}\left( \left( \begin{aligned} 0 \\ \boldsymbol{0} \end{aligned} \right),\left( \begin{matrix} \mathrm{Var}\left( g_{k} \right) & \mathrm{Cov}\left( g_{k},\mathbf{r} \right) \\ \mathrm{Cov}\left( \mathbf{r},g_{k} \right) & \mathbf{C} \end{matrix} \right) \right)$$

By standard multivariate normal conditioning,

$$g_{k}|\mathbf{y}_{\mathcal{O}}\boldsymbol{=}g_{k}|\boldsymbol{r\sim}\mathcal{N}\left( \mathrm{Cov}\left( g_{k},\mathbf{r} \right)\mathbf{C}^{-1}\mathbf{r},\mathrm{Var}\left( g_{k} \right)-\mathrm{Cov}\left( g_{k},\mathbf{r} \right)\mathbf{C}^{-1}\mathrm{Cov}\left( \mathbf{r},g_{k} \right) \right)$$

Thus, from standard multivariate normal conditioning, the posterior means of the three random effects for pair *k* are given by $E(u_{k}^{A}|\mathbf{y}_{\mathcal{O}})=\mathbf{k}_{k\mathcal{O}}^{A}\sigma^{2\left( A \right)}\mathbf{C}^{-1}\mathbf{r}$, ${E(u}_{k}^{B}|\mathbf{y}_{\mathcal{O}})=\mathbf{k}_{k\mathcal{O}}^{B}\sigma^{2\left( B \right)}\mathbf{C}^{-1}\mathbf{r}$, $E(u_{k}^{A\times B}|\mathbf{y}_{\mathcal{O}})=\mathbf{k}_{k\mathcal{O}}^{A\times B}\sigma^{2\left( A\times B \right)}\mathbf{C}^{-1}\mathbf{r}$.

Thus, the posterior mean of the total genotypic value for pair $k$ is $g_{k}=u_{k}^{A}+u_{k}^{B}+u_{k}^{A\times B}$.

Applying this separately to crops A and B, the posterior predictive mean of the Bayesian optimization objective function for pair $k$ at cycle $t$ is

$$\mu_{k,t}=g_{k}^{A}+g_{k}^{B}$$

The posterior variances of the three random effects are

$$\mathrm{Var}\left( u_{k}^{A}|\mathbf{y}_{\mathcal{O}} \right)=\mathbf{K}_{kk}^{A}\sigma^{2\left( A \right)}-\sigma^{4\left( A \right)}\mathbf{k}_{k\mathcal{O}}^{A}\mathbf{C}^{-1}\mathbf{k}_{\mathcal{O}k}^{A}$$

$$\mathrm{Var}\left( u_{k}^{B}|\mathbf{y}_{\mathcal{O}} \right)=\mathbf{K}_{kk}^{B}\sigma^{2\left( B \right)}-\sigma^{4\left( B \right)}\mathbf{k}_{k\mathcal{O}}^{B}\mathbf{C}^{-1}\mathbf{k}_{\mathcal{O}k}^{B}$$

$$\mathrm{Var}\left( u_{k}^{A\times B}|\mathbf{y}_{\mathcal{O}} \right)=\mathbf{K}_{kk}^{A\times B}\sigma^{2\left( A\times B \right)}-\sigma^{4\left( A\times B \right)}\mathbf{k}_{k\mathcal{O}}^{A\times B}\mathbf{C}^{-1}\mathbf{k}_{\mathcal{O}k}^{A\times B}$$

Because the three random effects are conditionally correlated given the observed data, the corresponding posterior covariances are

$$\mathrm{Cov}\left( u_{k}^{A},u_{k}^{B}|\mathbf{y}_{\mathcal{O}} \right)=-\sigma^{2\left( A \right)}\sigma^{2\left( B \right)}\mathbf{k}_{k\mathcal{O}}^{A}\mathbf{C}^{-1}\mathbf{k}_{\mathcal{O}k}^{B}$$

$$\mathrm{Cov}\left( u_{k}^{A},u_{k}^{A\times B}|\mathbf{y}_{\mathcal{O}} \right)=-\sigma^{2\left( A \right)}\sigma^{2\left( A\times B \right)}\mathbf{k}_{k\mathcal{O}}^{A}\mathbf{C}^{-1}\mathbf{k}_{\mathcal{O}k}^{A\times B}$$

$$\mathrm{Cov}\left( u_{k}^{B},u_{k}^{A\times B}|\mathbf{y}_{\mathcal{O}} \right)=-\sigma^{2\left( B \right)}\sigma^{2\left( A\times B \right)}\mathbf{k}_{k\mathcal{O}}^{B}\mathbf{C}^{-1}\mathbf{k}_{\mathcal{O}k}^{A\times B}$$

Therefore, the posterior variance of the total genotypic value for pair $k$ is

$$\mathrm{Var}\left( g_{k}|\mathbf{y}_{\mathcal{O}} \right)=\mathrm{Var}\left( u_{k}^{A}|\mathbf{y}_{\mathcal{O}} \right)+\mathrm{Var}\left( u_{k}^{B}|\mathbf{y}_{\mathcal{O}} \right)+\mathrm{Var}\left( u_{k}^{A\times B}|\mathbf{y}_{\mathcal{O}} \right)+2\mathrm{Cov}\left( u_{k}^{A},u_{k}^{B}|\mathbf{y}_{\mathcal{O}} \right)+2\mathrm{Cov}\left( u_{k}^{A},u_{k}^{A\times B}|\mathbf{y}_{\mathcal{O}} \right)+2\mathrm{Cov}\left( u_{k}^{B},u_{k}^{A\times B}|\mathbf{y}_{\mathcal{O}} \right)$$

Equivalently,

$$\mathrm{Var}\left( g_{k}|\mathbf{y}_{\mathcal{O}} \right)=\mathrm{Var}\left( g_{k} \right)-\mathrm{Cov}\left( g_{k},\mathbf{r} \right)\mathbf{C}^{-1}\mathrm{Cov}\left( \mathbf{r},g_{k} \right)$$

Applying this separately to crops A and B, and assuming independence between the crop-specific prediction models, the posterior variance of the objective function is given by

$$\sigma_{k,t}^{2}=\mathrm{Var}\left( g_{k}^{A}|\mathbf{y}_{\mathcal{O}}^{A} \right)+\mathrm{Var}\left( g_{k}^{B}|\mathbf{y}_{\mathcal{O}}^{B} \right)$$
